# Light-inducible T cell engagers trigger, tune and shape the activation of primary T cells

**DOI:** 10.1101/2022.04.15.488452

**Authors:** Morgane Jaeger, Amandine Anastasio, Sophie Brustlein, Renaud Vincentelli, Fabien Durbesson, Rémy Char, Maud Boussand, Mathias Lechelon, Rafael J. Argüello, Didier Marguet, Hai-Tao He, Rémi Lasserre

## Abstract

To mount appropriate responses, T cells integrate complex sequences of receptor stimuli perceived during transient interactions with antigen presenting cells. Although it has been hypothesized that the dynamics of these interactions influence the outcome of T cell activation, methodological limitations have hindered its formal demonstration. Here, we have engineered the Light-inducible T cell engager (LiTe) system, a recombinant optogenetics-based molecular tool targeting the T Cell Receptor (TCR). The LiTe system constitutes a reversible molecular switch displaying exquisite reactivity. As proof of concept, we dissect how specific temporal patterns of TCR stimulation shape T cell activation patterns. We established that CD4+ T cells respond to intermittent TCR stimulation more efficiently than their CD8+ T cells counterparts and provide evidence that distinct sequences of TCR stimulation encode different cytokine programs. Finally, we show that the LiTe system could be exploited to create light-activated bispecific T cell engagers and manipulate tumor cell killing. Overall, the LiTe system provides new opportunities to understand how T cells integrate TCR stimulations and to trigger T cell cytotoxicity with a high spatiotemporal control.

## INTRODUCTION

In addition to the nature and intensity of a stimulus, its dynamic features (frequency, duration) also constitute a layer of information that is decoded by cells and translated into specific outcomes (Purvis and Lahav, 2013). *In vivo*, T cell activation is usually initiated through transient and sequential interactions with different antigen presenting cells (APCs) presenting cognate antigenic peptide-MHC complex (pMHC) (Hugues et al., 2004; Mempel et al., 2004; Moreau and Bousso, 2014). At each interaction, the signaling of the T Cell Receptor (TCR) is triggered (Friedman et al., 2010) and T cells appear able to integrate, or sum, the signals perceived during series of TCR stimulations (Clark et al., 2011; Faroudi et al., 2003; Harris et al., 2021; O’Donoghue et al., 2021). The temporal pattern of TCR stimulation is thought to influence the outcome of the T cell activation (Hugues et al., 2004; Marangoni et al., 2013), although no causal relation has yet been formally demonstrated. A lack of methods enabling an accurate and reversible spatiotemporal manipulation of primary T cell stimulation has, to date, hindered our understanding of the cellular mechanisms behind the integration of activation signals and of the influence of signal dynamics on activation outcome.

Optogenetics is a technology based on the use of light-responsive protein domains from plants, algae or prokaryotes, to confer light-responsive capacity to targeted biological processes (Tan et al., 2022). The spatiotemporal control of cellular activities permitted with light has presented new opportunities in fundamental and biomedical research. Recently, optogenetic systems have been implemented in T cell models such as T cell lines or transduced pre-activated T cells, including chimeric antigen receptor T cells (CAR T cells) (Bohineust et al., 2020; Huang et al., 2020; O’Donoghue et al., 2021; Tischer and Weiner, 2019; Yousefi et al., 2019; Zhao et al., 2019). These elegant methodologies offered a new capacity to manipulate T cells and provided insight into the mechanism of T cell activation. However, they rely on the transduction into cells of genes encoding modified light-responsive proteins, preventing to study primary T cells that are known to be resistant to genetic modifications without previous *in vitro* pre-activation.

In order to determine the influence of TCR signal dynamics on primary T cell activation, we developed the LiTe system, an original optogenetic recombinant tool to control in space and time TCR stimulation using light. We demonstrated that LiTe system allows for accurate, tunable and reversible TCR stimulation, and can efficiently induces T cell effector functions. This molecular system showed an exquisite reactivity to photostimulations, is fully reversible and displayed a remarkable robustness as it keeps its functionality over days. Using specific temporal pattern of TCR stimulation, we were able to differentially activate the CD4^+^ and the CD8^+^ T cell subsets and to influence the outcome of the T cell compartment activation programme. Therefore, we formally demonstrate here the causal relationship linking TCR stimulation dynamics and the outcome of the T cell activation. Finally, we extended the LiTe system concept to generate the LiTe-Me system, a light-inducible bispecific T cell engager that enables light-targeted melanoma cell killing, *in vitro*.

## RESULTS AND DISCUSSION

### Design and validation of the LiTe system, a novel optogenetics-based recombinant molecular system for reversible control of TCR stimulations with light

We engineered the LiTe system, a recombinant and reversible light-inducible stimulatory system to control the activation of untouched primary T cells. Its rationale relies on the fact that a non-agonist monovalent Fab can become a potent agonist upon oligomerization (Mijares et al., 2000). The LiTe system has two components: (i) the N-terminal phytochrome B_1-651_ region from *A. thaliana* linked to the phycocyanobilin (PhyB) and (ii), the LiTe protein that combines the phytochrome interacting factor 6 (PIF6) to the Fab fragment from the H57 mAb, an agonist targeting the mouse TCR β-chain (Fig. 1a and Supplementary Fig. 1). Under red light illumination (e.g. 656 nm), PhyB captures the PIF6 domain of the LiTe protein. Far-red illumination abrogates this interaction. Once the LiTe proteins bind TCRs, their light-controlled capture by PhyB-coated supports induces reversible TCR triggering (Fig. 1b). We validated the ability of the LiTe protein to bind specifically to the TCR (Supplementary Fig. 2a,b). Next, we verified that its interaction with PhyB occurred only under red light exposure (Supplementary Fig. 2c,d).

**Figure 1.**
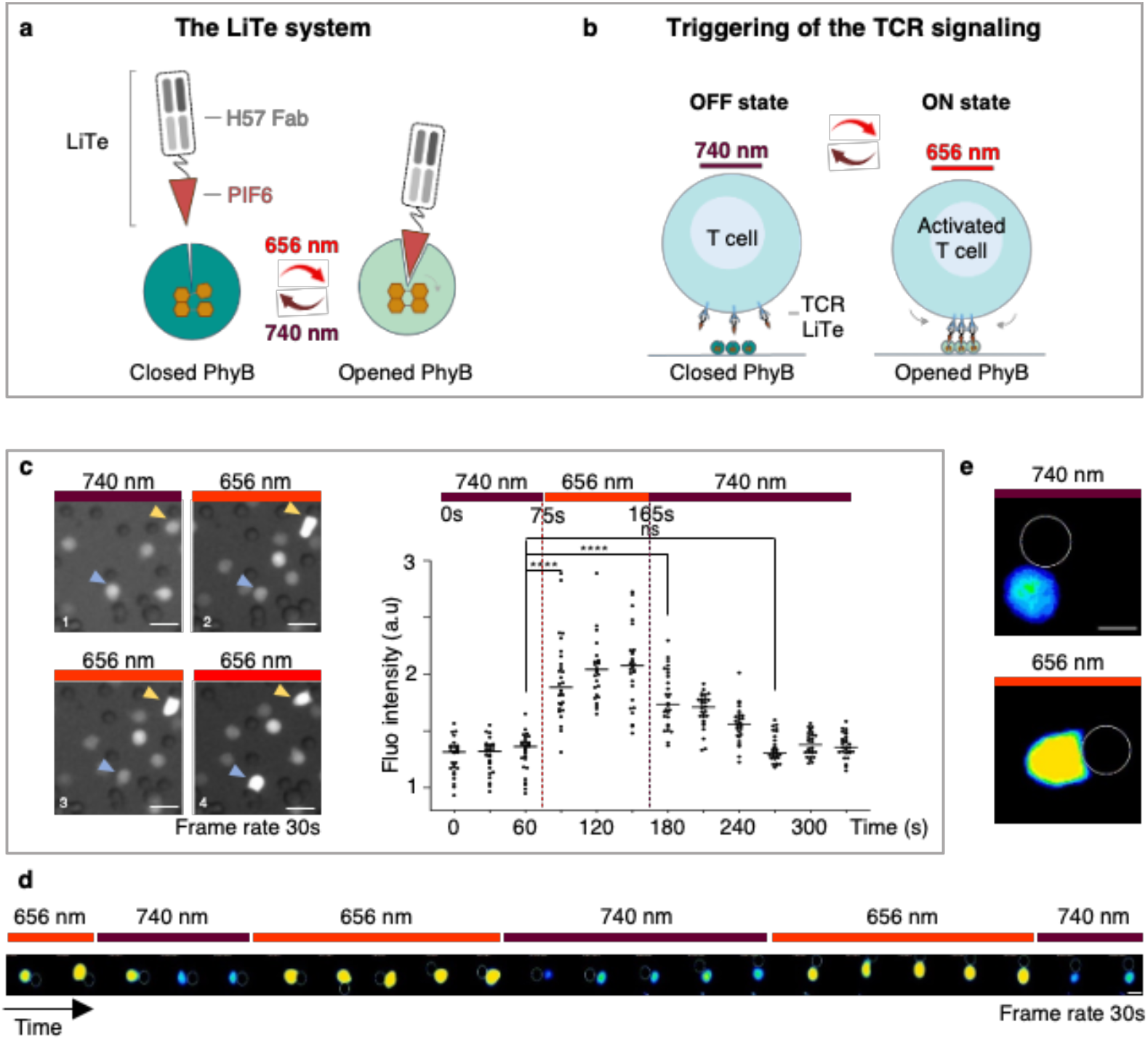
Design of the LiTe system for reversible triggering of TCR signaling in response to light. **a**, The LiTe system is in two parts: LiTe protein, a recombinant protein linking the PIF6 domain to the anti-mouse TCR H57-597 Fab and PhyB_1-651_. Exposure of the LiTe system to red light opens the PhyB binding site for PIF6 whereas far-red light reverses this conformational change. **b**, Exposure of mouse T cells in the presence of the LiTe system to 656 nm light induces the binding of TCR-bound LiTe proteins to PhyB-coated surface, leading to TCR aggregation/immobilization and triggering. This process is reversed by 730-760 nm light. **c**, Primary T lymphocytes were loaded with PBX, a calcium sensitive dye, then incubated with the LiTe protein before being mixed with PhyB-coated beads and imaged by fluorescence microscopy at 37°C. Left panel, time-lapse images of the cells exposed to 740 nm light, then to 656 nm light in the presence of the LiTe system. Arrows indicate cells in contact with PhyB-coated beads over time. Right panel, quantification of relative fluorescence intensity of cells in contact with PhyB-coated beads and exposed to the specified light sources (n = 30 cells, representative of >3 experiments; **** P < 0.0001, Wilcoxon-Mann-Whitney test; scale bar: 10 μm). **d**, Time-lapse images of a T cell under iterative stimulation/resting cycles (scale bar: 5 μm, frame rate: 30 s). **e**, Morphological changes of a lymphocyte in contact with PhyB-coated bead and in response to exposure to 740 nm (left) or 656 nm (right) lights; scale bar: 5 μm.

We evaluated the performance of the LiTe system on living primary T cells by recording its ability to trigger intracellular calcium influx as an early, sensitive and dynamic readout of TCR engagement (Harris et al., 2021; Huse et al., 2007; Varma et al., 2006). Primary CD8^+^ T cells were loaded with PBX calcium-sensitive dye and the LiTe protein before incubation with PhyB-coated beads; they were then imaged with a videomicroscope at 37°C. Switching light from 740 nm to 656 nm triggered an increase in intracellular calcium within tens of seconds solely for T cells in contact with PhyB-coated beads (Fig. 1c and Supplementary Fig. 3). The calcium concentration gradually returned to the basal level within two minutes after switching the light back to 740 nm. These dynamics are consistent with previous observations for TCR-induced calcium influx during antigen recognition engagement (Harris et al., 2021; Huse et al., 2007; Varma et al., 2006). Moreover, the LiTe system enabled the delivery of pulsed stimuli to T cells via the application of iterative cycles of 656/740 nm light exposure, as shown by the synchronization of intracellular calcium fluxes with the light cycle dynamics (Fig. 1d). Also, under 656 nm light, the shape of T cells in contact with PhyB-coated beads was reminiscent of that reported at the immunological synapse; this suggests the engagement of a large number of TCRs (Fig. 1e). Altogether, these observations demonstrate that the LiTe system has the unique ability to operate on primary T cells without requiring their genetic manipulation. It constitutes an ON/OFF molecular switch enabling precise spatiotemporal control of T cell activation via strong, reversible and iterative TCR stimulation.

### LiTe system triggers T cell effector functions and lay the fundation for light-inducible bispecific T cell Engagers for tumor cell killing

We next wanted to determine whether the LiTe system could induce functional activation of primary T cells. Activated T cells are characterized by the surface expression of activation markers, as well as by the onset of effector functions such as cytokine secretion or cytotoxicity. Thus, primary CD8 T cells were incubated with the LiTe system, in the presence of soluble anti-CD28 mAbs that increased the fraction of responding cells (Supplementary Fig. 4). Cells were placed on an illumination device (optoPlate-96, (Bugaj and Lim, 2019)) and illuminated for 18 hours with light at 630 nm (ON), or 780 nm (OFF). Next, T cell activation was assessed by flow cytometric analysis of late activation markers (Fig. 2a). Under 780 nm light conditions, the cells remained in resting state. In contrast, the 630 nm light triggered potent T cell activation, as observed by an increase in CD69 and CD25 expression, and a decrease in CD62-L. Under these conditions, up to 80% T cells were activated such as with anti-CD3/anti-CD28 stimulation. As expected, neither the 630 nm light nor LiTe system alone could induce any T cell activation (Supplementary Fig. 5a). Also, no phototoxicity was noted with the low light power used in these experiments (0.14 mW.cm^-2^ at 630 nm and 2.8 mW.cm^-2^ at 780 nm; Supplementary Fig. 5b). We then determined whether the LiTe system permits the tuning of T cell activation using combined illuminations at different ratios of 630 nm/780 nm (Fig. 2b). Flow cytometry analysis of CD69 and CD25 upregulation revealed a proportional relationship between T cell activation and the ratio of 630 nm to 780 nm light power. Therefore, the LiTe system allowed adjustable control of the extent of T cell activation.

**Figure 2.**
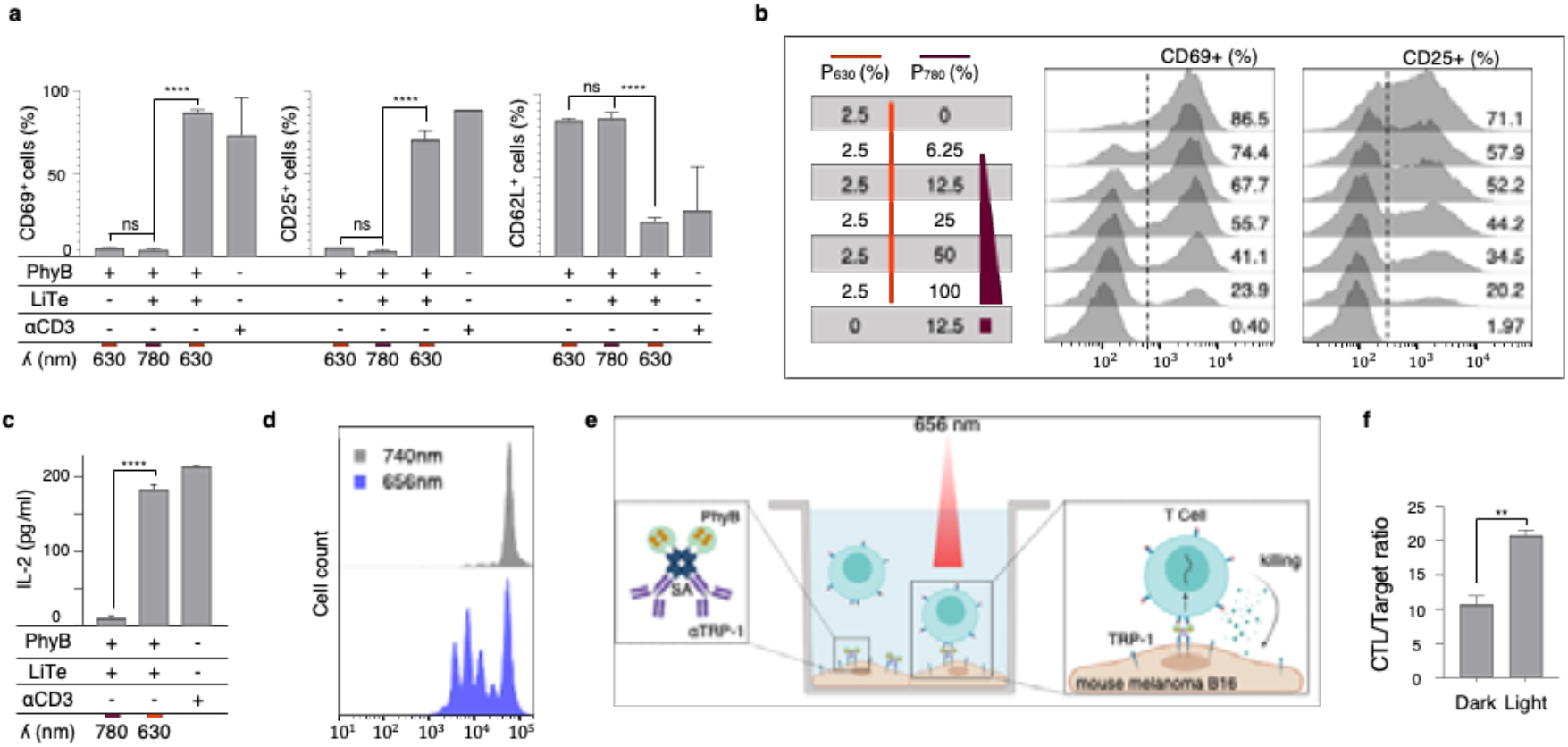
LiTe-driven TCR photostimulation is tunable and leads to effective T cell activation. **a**, Primary T cells were incubated with the LiTe system and anti-CD28 antibody, then illuminated 18 h in optoPlate at the specified wavelength. Flow cytometry analysis of CD69, CD25 or CD62L positive cells in response to 630 or 780 nm light (n > 3, mean +/-SEM are shown; *** P < 0.001, **** P < 0.0001, Student t-test). **b**, Same experimental conditions as in **a**, except that cells were exposed to different ratios of 630 nm/780 nm light power (expressed as % of max power intensity) and analyzed for the expression of CD69 or CD25 by flow cytometry. **c**, Same experimental conditions as in **a**, except that IL-2 secretion was analyzed in the cell supernatant by ELISA (n = 3; mean +/-SD are shown; **** P < 0.0001, Student t-test). **d**, T Cells treated as in **a** were illuminated 72 h with 780 nm light (upper panel) or 630 nm light (lower panel) before the analysis of their proliferation by measurement of CellTrace Violet dilution by flow cytometry (representative of n > 3 experiments). **e**, LiTe-Me system design. PhyB is combined to streptavidin (SA) and to the melanoma cell-targeting antibody TA-99 (specific for surface molecule TRP-1). When bound to melanoma cells, the Me-PhyB complex can engage LiTe-bound CTL when exposed to 656 nm light, and trigger tumor cell killing. **f**, CTLs were dropped on a B16F10 melanoma cell monolayer in the presence of the LiTe-Me system, and illuminated or not with a 656 nm light for 18 h at 37°C. Flow cytometry analysis of the CTL/target cell ratio, identified respectively with anti-CD45 and anti-TRP-1 mAbs (n = 2; mean +/-SD are shown; ** P < 0.001, Student t-test).

To assess the ability of the LiTe system to induce T cell effector functions, we first monitored cytokine production. Purified primary CD8^+^ T cells were stimulated as above with 630 nm light and secreted IL-2 was quantified by ELISA (Fig. 2c). The LiTe system appeared very effective at inducing IL-2 production by the T cells, only when exposed to 630 nm light. Next, to further characterize the LiTe-induced activation, the proliferation of T cells treated as above and exposed to 630 nm or 780 nm light for 72 hours was analyzed by flow cytometry (Fig. 2d). We observed that the LiTe system induces a strong proliferation of the T cells only under 630 nm light exposure. Overall, this demonstrates the ability of the LiTe system to induce full T cell activation. It also reveals that this recombinant molecular system is capable of delivering long-term stimulations.

A fundamental effector function of the CD8^+^ T cell compartment is the killing of pathological cells, such as cancer cells. We therefore wished to evaluate the capacity of the LiTe system to trigger tumor cell killing by syngenic CD8^+^ effector T cells. To this end, we modified the LiTe system to generate the LiTe-Me system in which PhyB is targeted to the surface of <u>me</u>lanoma cells by coupling to a mAb specific for TRP-1, a melanoma-associated antigen (Fig. 2e). In practice, we generated a molecular complex containing streptavidin and biotinylated forms of PhyB and anti-TRP1 mAb (Supplementary Fig. 6). Then, CD8^+^ effector T cells and the LiTe-Me system were dropped on a B16F10 melanoma cells monolayer and illuminated or not with red light for 18 h. Flow cytometry measurement of T cell/B16F10 ratios showed a 2-fold increase only when the cells were in the presence of the LiTe-Me system and illuminated with red light (Fig. 2f).

Altogether, these experiments show that the LiTe system enables specific and adjustable activation of primary T cells, leading to the establishment of their effector functions. The LiTe-Me system provides control of T cell activity at the cell/cell interface, without the use of beads or any artificial surfaces. More importantly, we provide *in vitro* evidence that LiTe-Me acts as a light-inducible Bispecific T cell Engager (BiTE), allowing spatio-temporal control with light of tumor cell killing by CD8^+^ effector T cells.

### The temporal pattern of TCR stimulation shapes the activation outcome of the T cell compartment

Previous studies have suggested that the dynamics of T cell/APC interactions influences the outcome of T cell activation (Hugues et al., 2004; Marangoni et al., 2013). Yet, the causal relationship has not been formally demonstrated. With this in mind, we next wished to exploit the flexibility of the LiTe system to explore how the temporal pattern of the TCR stimulation influences the outcome of T cell population activation. We also aimed to determine whether the LiTe system could allow manipulation/orientation of the adaptive immune response.

We first confirmed that PhyB remained functional after hours of exposure to illumination cycles alternating between 630 nm and 780 nm light (Supplementary Fig. 7). Then, purified primary T cells were incubated with the LiTe system under different conditions of light exposure, and soluble anti-CD28 mAb. Cells were stimulated with time-gated 630 nm light exposure in continuous or pulsed mode, and then the CD69 and CD25 upregulation in CD8^+^ T and CD4^+^ T cell subsets was analyzed by flow cytometry (Fig. 3a). Upon continuous TCR stimulations, the fraction of cells upregulating CD69 and CD25 increased with the duration of 630 nm light exposure, to similar extents for the CD8^+^ and CD4^+^ subsets although the latter responded slightly better to short stimulations (Fig. 3b). However, when T cells were exposed to different temporal patterns of pulsed stimulation for 18h, CD4^+^ T cells clearly showed a higher response than CD8^+^ T cells (Fig. 3c). For the short 2 min ON/4 min OFF cycles, the proportion of activated cells was up to 2-fold higher in CD4^+^ than in CD8^+^ T cell subsets (57% versus 28% of CD69+ cells, respectively). In comparison to continuous stimulations, pulsed stimulations favored the activation of CD4^+^ T cells over CD8^+^ T cells, as illustrated by the ratios of activated CD4^+^/CD8^+^ T cells (Fig. 3d). Of note, the duty cycles were held constant (33%, 6 h of cumulated ON state over an 18-h period) between different pulse cycles that ranged from 6 to 90 min. The cycle length had only a moderate influence on primary T cell activation (Fig. 3c).

**Figure 3.**
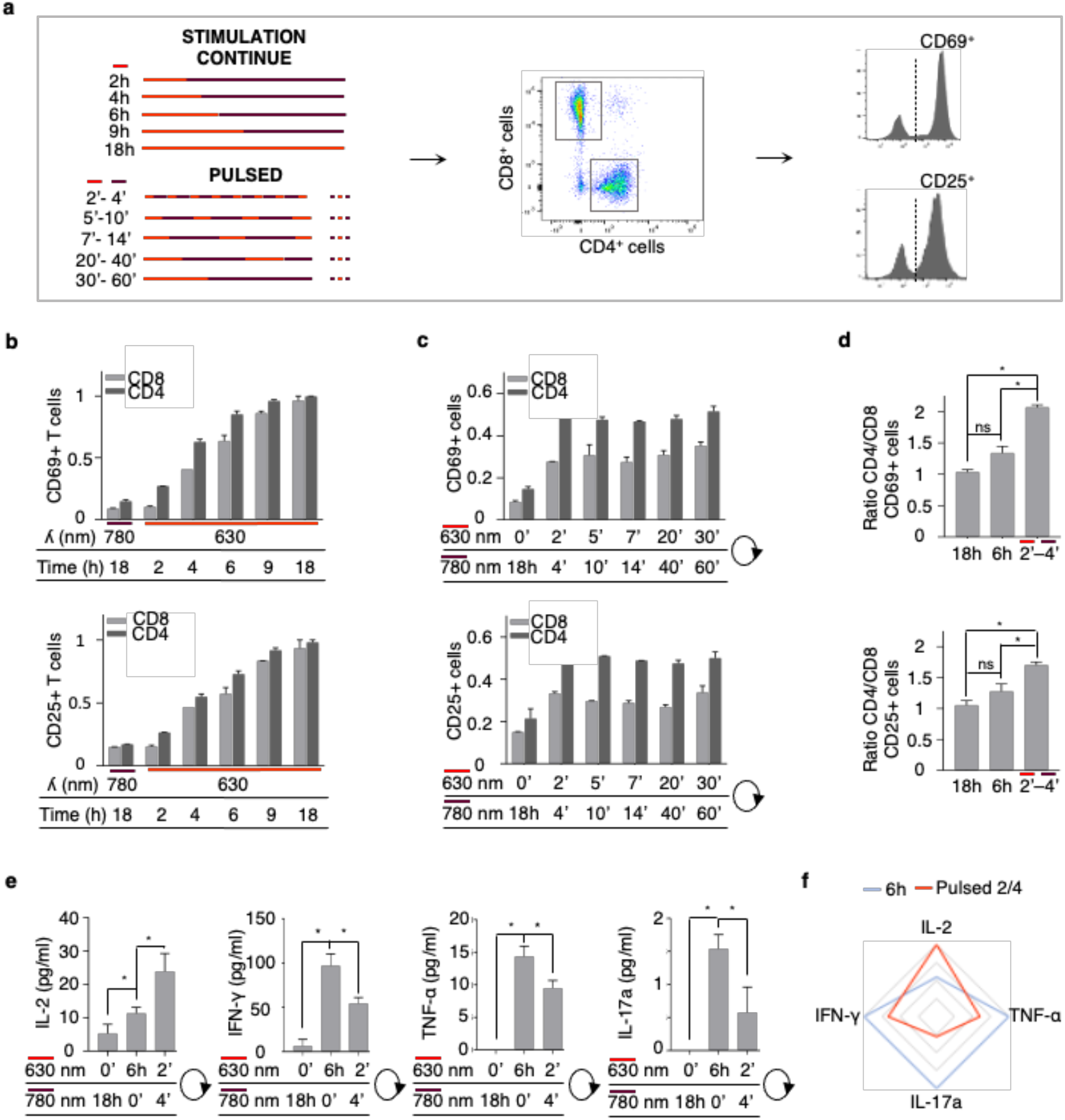
LiTe system enabled the manipulation of the adaptive immune response. **a**, Primary T cells were incubated with the LiTe system and anti-CD28 antibody, then illuminated for 18 h in optoPlate under continuous or pulsed illumination conditions. All pulsed illumination conditions correspond to 6 h of cumulative exposure time to 630 nm light. CD4^+^ and CD8^+^ T cell subsets were analyzed by flow cytometry for CD69 and CD25 expression. **b**, Flow cytometry analysis of the T cell activation following different periods of continuous photostimulation (n > 3; mean +/-SEM are shown; normalized data to 18 h continuous photostimulation value). **c**, same as in **b**, but for pulsed photostimulation of different durations (n > 3; mean +/-SEM; normalized data to 18 h continuous photostimulation value). **d**, Ratio of CD69^+^ CD4^+^/ CD69^+^ CD8^+^ T cells (upper panel) and CD25^+^ CD4^+^/ CD25^+^ CD8^+^ T cells (lower panel) under different photostimulation conditions. The 2’-4’ pulsed stimulation condition over 18 h corresponds to a cumulated 6 h exposure to the 630 nm light (n > 3; mean +/-SD are shown; data normalized to 18 h continuous photostimulation value, * P < 0.05, Wilcoxon-Mann-Whitney test). **e**, Measurement of cytokine secretion in the supernatant of T cells stimulated or not 6 h in continuous or pulsed modes. IL-2 has been detected by ELISA, IFN-γ, TNF-α and IL-17 by multiplex assay (n = 2; mean +/-SD; * P < 0.05, Wilcoxon-Mann-Whitney test). **f**, Polar representation of the cytokine panels produced by the T lymphocytes stimulated after the continuous (in blue) or the pulsed (in red) stimulations shown in **e** (normalized to the maximum value detected for each cytokine).

The differential responses shown by T cell subsets to pulses stimulations raised the possibility of using the LiTe system to qualitatively manipulate the activation of the T cell compartment. We therefore characterized the panel of cytokines produced by primary T cells following 6 h continuous or pulsed stimulations (iterative 2 min ON/4 min OFF). Interestingly, pulsed stimulations of T cells induced higher amounts of IL-2 production than continuous stimulations, but lower amounts of IFN-γ, TNF-α and IL-17 (Fig. 3e,f).

Altogether, these experiments revealed a remarkably enhanced ability of CD4^+^ T cells to integrate series of short-lived TCR stimulations, when compared to CD8^+^ T cells. They also demonstrated that TCR stimulation dynamics directly influences the panel of cytokines secreted by the T cell compartment. Finally, the combination of the LiTe system with specific illumination strategies allows therefore the manipulation of T cell compartment activation and the release of associated cytokines.

Optogenetics is a promising technology for manipulating cellular activity in fundamental and biomedical research. However, it has been limited, to some degree, by the constraint of genetically modifying the target cells, making difficult the study or manipulation of primary cells. Our results in primary T cells show that use of the fully recombinant LiTe system can overcome this limitation. By way of its modular design, the LiTe system concept can to be easily adapted to other biological systems. By providing the spatiotemporal control of TCR stimulation, the LiTe system enables both quantitative and qualitative manipulation of T cell activation. As mentioned above, nearly two decades ago it was proposed that the dynamics of T cell stimulation could qualitatively influence the cellular response (Hugues et al., 2004; Marangoni et al., 2013). We provide here direct evidence of a causal relationship linking the dynamics of TCR stimulation and the outcome of T cell activation. Further research is needed to refine our knowledge on this relationship and to determine the extent to which controlling the dynamics of TCR stimulation with the LiTe system would enable the selective induction of different T cell activation programmes (e.g. tolerance, TH-1, TH-2, or TH-17 responses…). In this context, Achar et al have recently reported that distinct cellular responses of the T cells are elicited by antigenic peptides of different affinity for the OT-I TCR (Achar et al., 2022). Peptide affinity influences both the half-life of TCR-pMHC binding, and the stability of T cell/APC interactions (Moreau et al., 2012). As the LiTe system, based on PhyB photoactivation, enables the control of the duration of TCR engagement at the molecular level (Yousefi et al., 2019), but also at the cellular level, it could provide new opportunities to uncover the molecular mechanisms underlying the induction of such a diversity of cellular responses.

With new technologies to illuminate optogenetic tools in tissue (Joshi et al., 2019; Nguyen et al., 2021; Bansal et al., 2022), optogenetics is now being applied in onco-immunology research and has enabled the production of light-inducible CAR T cells in mice (Huang et al., 2020; Zhao et al., 2019). In this context, the LiTe-Me system presented here has provided a proof-of-concept of the relevance of optogenetics to also control the bioactivity of BiTEs by light. BiTEs are promising novel therapies against cancer, but their usage is sometime limited by the induction of severe adverse effects due to an immune activation at the systemic level, which can be very deleterious and some time letal. Overcoming these adverse effect is a central challenge in onco-immunology (Young et al., 2018). Spatiotemporal control by light of the activity of light-responsive BiTEs, such as the LiTe-Me system, could be a relevant strategy to generate new highly targeted immunotherapies with decreased adversed effects. Furthermore, considering the ability of the LiTe system to shape the outcome of T-cell activation in response to specific time-gated photostimulations, light-controllable BiTEs could provide a new level of control of the BiTE-induced antitumor immune response. Indeed, differences in the production of IL-2, TNF-α, IL-17 and IFNγ in the tumor bed can have a strong influence on the recruitment and function of bystander T cells, NK cells, macrophages, neutrophils and other stromal cells. Future studies will be necessary to test LiTe-Me *in vivo* and determine the specific improvements needed to fully unleash its therapeutical potential. Its regulation with light in the red and the far-red regions, which penetrate tissues at the millimeter scale, constitutes a great advantage for these applications (Bansal et al., 2022).

Overall, the LiTe system constitutes an innovative and versatile molecular tool that allows precise spatio-temporal control of primary cells. It provided us the novel capacity to directly demonstrate the influence of the TCR stimulation dynamics on the outcome of the T cell activation. Moreover, the LiTe-Me system opens up new avenues in biomedical research for the development of future therapeutic approaches.

## MATERIALS AND METHODS

### Reagents and antibodies

Dulbecco’s modified Eagle medium (DMEM), Roswell Park Memorial Institute (RPMI) medium, Dulbecco’s Modified Eagle Medium/Nutrient Mixture F-12 (DMEM/F-12), sodium pyruvate (NaPy), L-glutamine (L-GLu), penicillin-streptomycin (Pen-Strep), geneticin, HEPES buffer solution, and β2-mercaptoethanol were purchased from Gibco. Phosphate-buffered saline (PBS), streptomycin sulfate salt, biotin, tris (2-carboxyethyl) phosphine (TCEP), Tween 20, and valproic acid sodium salt were supplied by Sigma Aldrich. Streptavidin-HRP was purchased from Beckman Coulter. Alexa Fluor 488 and caspase-3/7 activity detection dyes were purchased from Thermo Fisher, and Viobility™ 405/452 Fixable Dye from Miltenyi Biotec.

The monoclonal antibodies anti-CD28 (clone H37.51) and goat anti-mouse IgG-HRP were purchased from Thermo Fisher, APC anti-CD69 (clone REA937) and FITC anti-CD62L (clone MEL-14) from BioLegend, FITC anti-CD25 (clone 7D4) and anti-6xHis (clone F24-796) from BD Biosciences, AF647 anti-histidine (clone AD1.1.10) from Bio-RAD, and APC anti-CD45 (clone 30-F11) from eBioscience™. The TA-99 antibody (α-TRP1) was a gift from Innate Pharma SA. The anti-TCR β (clone H57-597) and anti-CD3ε (clone 145-2C11) were purified from hybridoma culture supernatants following standard protocols. H57-597 Fab fragment was prepared by papain digestion, purified by ion-exchange, and controlled by SDS-PAGE. Fluorescent labeling of proteins was performed according to the manufacturer’s instructions.

### Cell culture

The melanoma B16F10 cell line was cultured in the RPMI medium supplemented with 10% fetal bovine serum (FBS) with 7.5% CO_2_ at 37°C. The MCD4 cell line was derived from the mouse 3A9 CD4^+^ T cell hybridoma expressing a high TCR level specific for hen egg lysozyme (HEL) peptide bound to MHC II I-Ak molecules, as previously described (Sadoun et al., 2021). MCD4 cells were grown in RPMI medium supplemented with 5% FBS, 1 mM NaPy and 10 mM HEPES with 10% CO_2_ at 37°C. The human embryonic kidney 293T cell line (HEK293T) was grown in DMEM supplemented with 10% FBS, 1 mM NaPy, 2 mM L-Glu and geneticin with 7.5% CO_2_.

Primary T lymphocytes were isolated from lymph nodes of C57BL/6 mice using the Dynabeads™ Untouched™ mouse T cell kit (Invitrogen™) by negative selection. CD8^+^ T cells were isolated from lymph nodes of C57BL/6 Rag1^-/+^ OT-1^-/+^ mice and purified using the EasySep™ mouse CD8^+^ T cell isolation kit (STEMCELL Technologies) by negative selection. T cells were cultured in complete DMEM/F-12 medium (DMEM/F-12 supplemented with 10% FBS, 1 mM NaPy, 10 mM HEPES, 50 U/mL Pen-Strep, 0.05 mM β2-mercapto-ethanol) with 10% CO_2_ at 37°C.

The effector CD8^+^ T cells were cultured in 6-well plates at 0.625 10^6^ cells/mL in complete DMEM/F-F12 medium with 1 μg/mL anti-CD28 (clone H37.51). The 6-well plates were coated with 3 μg/mL anti-CD3ε (145-2C11) in PBS for 4 h at 37°C and washed three times with PBS before seeding the T cells. Cells were cultured for 48 h (37°C, 7.5% CO_2_), after which IL-2 (PeproTech) was added to a final concentration of 10 U/mL. The cells were then cultured for a further 48 h.

### Generation and production of PhyB

PhyB was produced in BL21(DE3) *E*.*coli* (Sigma Aldrich) transfected with three plasmids obtained from the laboratory of Dr K Rosen and Leung (Leung et al., 2008) that encode the heme oxygenase 1 (Ho1), phytochromobilin (PΦB) synthase lacking the transit peptide (ΔHY2), and Phytochrome B (PhyB_1-651_). The latter was modified to add a Strep-tag at the C terminus of PhyB_1-651_. Alternatively, we used the plasmid pMH1105 (Horner et al., 2020), a gift from Dr Wilfried Weber (Addgene, plasmid # 131864 ; http://n2t.net/addgene:131864 ; RRID:Addgene_131864) to produce a biotinylated form of PhyB; this plasmid encodes the PhyB_1-651_, HO1 and PCB:ferredoxin oxidoreductase (PcyA). For the production of PhyB, the transfected bacteria were grown at 30°C in lysogeny broth medium supplemented with the selection antibiotics, then induced with 1 mM Isopropyl-β-D-1-thiogalactopyranoside (IPTG) at OD 600 nm between 0.6 and 0.8 and grown overnight at 18°C in the dark at 150 rpm. For biotinylated PhyB production, biotin was added (5 μM) to the culture medium allowing the biotinylation of the Avi-Tag. Bacteria were harvested by centrifugation for 8 min at 6,500 g and 4°C, then resuspended in lysis buffer (50 mM HEPES, 500 mM NaCl, 5% glycerol, 0.5 mM TCEP, 20 mM imidazole, pH 7.4) and shock frozen in liquid nitrogen before being stored at -80°C. Bacteria were disrupted by adding DNAse (10 μg/mL) and MgSO_4_ (20 mM) during 20 min and were then sonicated. Lysate was cleared from debris by centrifugation at 30,000 g and 4°C for 30 min before being loaded onto a Ni-NTA Superflow cartridge (Qiagen) using an Äkta Explorer chromatography system (GE Healthcare), then purified. The purified protein buffer was exchanged to PBS containing 0.5 mM TCEP using a HiPrep 26/10 desalting column (GE Healthcare).

### Generation and production of the LiTe

In order to produce the LiTe protein, gene encoding the H57 Fab Light Chain (LC) was cloned into pTT22 eukaryotic expression vector (pTT22-H57Fab-LC plasmid), and the H57 Fab Heavy Chain (HC) linked to the PIF6 domain with a 6xHis tag (Supplementary Fig. 1) within the pYD7 eukaryotic expression vector (pYD7-H57Fab-HC-PIF plasmid). All cloning reactions were performed using In-Fusion HD cloning kit (Clontech) in a standard reaction mixture.

The HEK293T cell line was then transfected with both the pYD7-H57Fab-HC-PIF plasmid and the pTT22-H57Fab-LC plasmid (ratio 1:3) using polyethylenimine (PEI, Polysciences). Cells were then maintained in DMEM supplemented with 2% FBS, 0.5% Tryptone TN1 (OrganoTechnie), 1.25 mM valproic acid and geneticin, at 37°C with 5% CO2. Supernatants were harvested 7 days later and the protein purified by Ni-NTA affinity chromatography. The purified protein buffer was exchanged to PBS using a Slide-A-Lyzer™ Dialysis Cassette (Thermo Fisher).

### Western blot analysis

Purified proteins in reduced 5X Laemmli loading buffer were denatured for 5 min at 95°C before being loaded onto a 10% SDS–PAGE gel, separated by electrophoresis and transferred onto nitrocellulose membrane (LI-COR). Membranes were then blocked with 4% bovine serum albumin (BSA) in TBS/T buffer (137 mM NaCl, 20 mM Tris pH 7.6, 0.1% Tween 20). Proteins were detected with reagents coupled to HRP and revealed by chemiluminescence with the Pierce™ ECL Western kit (Thermo Scientific™) and Azure biosystems 300 imaging system.

### Flow cytometry analysis

Cells were washed twice with FACS buffer (PBS, 0.5% FBS), then incubated for 45 min at 10°C before being labeled with antibodies in FACS buffer. After 3 washes, cells were resuspended in FACS buffer and analyzed by flow cytometry on a BD FACSCanto™. Data were analyzed with FlowJo software (Treestar Inc., CA).

### Spectral analysis of PhyB photoswitching

Spectral analysis of PhyB photoswitching was measured with a NanoDrop™ One (Thermo Scientific™). The absorption spectrum of PhyB was measured after 3 min under light exposure at 680 nm. The samples were then exposed to 740 nm light for 1 h before recording a new absorption spectrum.

### Light-controlled PhyB/LiTe pull-down assay

Streptavidin-coated magnetic beads (MagStrep “type3” XT beads 5% suspension, Iba) were washed three times in PBS supplemented with 0.5 mM TCEP with a magnetic separator (Iba). The beads were then incubated with 20 μg/mL of PhyB for 1 h at 4°C, then washed 3 times before adding the LiTe protein at a concentration of 20 μg/mL. The mix samples were illuminated 10 min with light at 656 nm or 740 nm. After 3 washes, the samples were boiled and the eluted proteins analyzed by western blotting.

### Photostimulations and calcium imaging

A 20 μg/mL solution of PhyB protein was added to washed streptavidin-coated polystyrene beads (5.0-5.9 μm, Spherotech), before a 2h incubation on ice. After washing in PBS supplemented with 0.5 mM TCEP, the beads were resuspended in complete DMEM/F-12 medium without phenol-red and distributed in Lab-Tek™ II Chambered Coverglass (Thermo Fisher). Primary T lymphocytes were loaded with PBX calcium reporter according to the manufacturer’s instructions, then washed before incubation with 20 μg/mL of LiTe protein for 45 min at room temperature, and finally washed again before incubation with PhyB-coated beads. The samples were illuminated with a 656nm or 740nm LED system (Mightex) and imaged on a videomicroscope, at 37°C.

### T cell activation analyses

A 20 μg/mL solution of PhyB protein was added to washed streptavidin-coated polystyrene beads and then incubated for 1 h in ice. After washing with PBS supplemented with 0.5 mM TCEP, the beads were resuspended in complete DMEM/F-12 medium. The beads were distributed in black clear-bottom 96-well plates (Greiner). Plates were illuminated on optoPlate for 3 min under 780 nm light to ensure a closed conformation of all PhyB proteins. Primary T cells labeled with 0.3 μg/mL of LiTe protein were added at a concentration of 5 10^5^ cells/well in the presence of 2 μg/mL anti-CD28 antibody. The plates were incubated on the optoPlate located in the cell culture incubator, and exposed to the indicated photostimulation programs. The harvested T cells were then analyzed by flow cytometry to measure cell surface protein expression, and by ELISA of their supernatants to measure Il-2 secretion. Plates were read at 450 nm with a Tecan INFINITE M1000 PRO. For T cell proliferation analysis, T cell were labeled with the Celltrace Violet Cell Proliferation Kit (Thermo Fischer Scientific) 20 min à 37°C, before beeing stimulated. After 72h of illumination at 630 nm or 780 nm, the Celltrace Violet dilution was analyzed by flow cytometry.

### Melanoma-targeted PhyB generation and enrichment

Biotinylated PhyB was added to a streptavidin solution in PBS/TCEP with 2:1 molar ratio under constant vortexing before the addition of an excess of biotinylated TA-99 antibody. Samples were separated by HPLC, and the protein content evaluated by absorbance at 280 nm and 680 nm for total protein and PhyB detection, respectively. The collected fractions were then analyzed by western blotting and revealed with streptavidin-HRP to detect biotinylated proteins.

### Measurement of light-induced tumor cell killing by flow cytometry

A total of 2.10^5^ melanoma B16F10 cells per well were plated in a 96-well plate and loaded with the melanoma-targeted PhyB. Effector CD8^+^ T cells were cocultured at 10:1 ratio with the target cells in a total volume of 200 μl/well of RPMI supplemented with 10% FBS, 1 mM NaPy, 10 mM HEPES, 0.05 mM β2-mercapto-ethanol and the LiTe protein (0.3 μg/mL). Cells were illuminated with the indicated light for 18 h at 37°C. Then, the CTL/target cells ratio, identified respectively with an anti-CD45 and an anti-TRP1 mAb, were determined by flow cytometry.

### Statistical analyses

Quantitative data were reported as means ± SD or ± SEM. Excel (Microsoft) and Prism (GraphPad Software) were used for statistical analyses. The non-parametric Wilcoxon-Mann-Whitney test or the Student t test were used for the evaluation of significance, as indicated in the figures. The sample number (*N*) refers to the number of biological replicates. n.s. = not significant (*P* > 0.05), \**P* < 0.05, \*\**P* < 0.01, \*\*\**P* < 0.001 and \*\*\*\**P* < 0.0001.

## ACKNOWLEDGMENTS

We thank J.M. Ingargiola from the Institut des Sciences du Mouvement and Dr. S. Mailfert from the PICSL imaging facility of the CIML (ImagImm), for the construction of the optoPlate, Drs Leung and Rosen for plasmids encoding PhyB and related enzymes, Drs Horner and Weber for advice and plasmid for PhyB production, A. Formisano for AF488-H57-597 Fab generation and coupling, Dr L. Gauthier for TA-99 antibody supply and advice, C. Debarnot for advice in protein production, and Dr B. Malissen for advice. We thank the CIML cytometry platforms for their technical and methodological support. We acknowledge the PICSL imaging facility of the CIML (ImagImm), member of the national infrastructure France-BioImaging supported by the French National Research Agency (ANR-10-INBS-04). **Funding**: This work was supported by institutional funding from the French National Institute of Health and Medical Research (Inserm), Centre National de la Recherche Scientifique (CNRS), and Aix-Marseille-Université (AMU), and program grants from the French National Research Agency (ANR-14-CE14-0014- 01 to R.L.; ANR-18-CE15-0021-02 to D.M., ANR-17-CE15-0032 to H.T.H) and SATT Sud-Est (SATT N°191702 to D.M.). M. J. was funded by fellowship from INSERM/ Région SUD/Innate Pharma (RSE18003AAA), and by the French Foundation for Medical Research (FRM: FDT202106013223).

## Author contribution

M.J, A.A, S.B, R.L designed the study, performed the experiments and analyzed the results. F.D, R.V, R.C, R.J.A, M.B, M.L assisted in the realization and interpretation of the experiments. R.L supervised and direct the research. R.L, H-T.H, D.M participate in the acquisition of the financial supports of the project. R.L, D.M, M.J wrote the manuscript. All authors discussed the results and contributed to editing of the manuscript. The manuscript has been revised for the English by Angloscribe, an independent scientific language editing service. Competing interests: The authors declare no competing interests. Patent application PCT/EP2019/076914 and EP22305545.0 corresponding respectively to the LiTe and the LiTe-Me systems have been filed.

## Data and materials availability

All data needed to evaluate the conclusions in the paper are present in the paper and/or Supplementary Materials. Additional data related to this paper may be requested from the authors.

