## Supplemental figures for "Light-inducible T cell engagers trigger, tune and shape the activation of primary T cells"

### Supplementary Figures

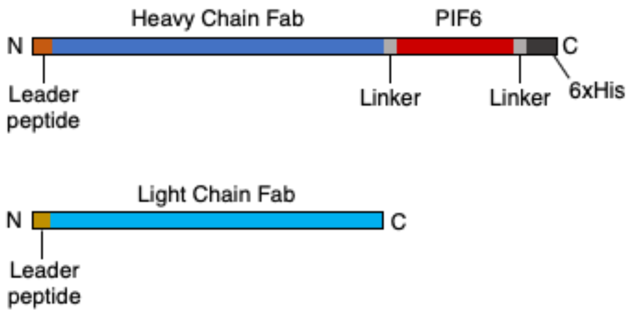


Protein of the Heavy Chain Fab - PIF :
MEFGLSWVFLVALFRGVQCEVYLVESGGDLVQPGSSLKV
SCAASGFTFSDFWMYWVRQAPGKGLEWVGRIKNIPNNY
ATEYADSVRGRFTISRDDSRNSIYLQMNRLRVDDTAIYYC
TRAGRFDHFDYWGQGTMVTVSSASTKGPSVFPLAPSSKS
TSGGTAALGCLVKDYFPEPVTVSWNSGALTSGVHTFPAV
LQSSGLYSLSSVVTVPSSSLGTQTYICNVNHKPSNTKVDK
RVEPKSCDKTGAGSGSGSGSGSMMFLPTDYCCRLSDQE
YMELVFENGQILAKGQRSNVSLHNQRTKSIMDLYEAEYNE
DFMKSIIHGGGGAITNLGDTQVVPQSHVAAAHETNMLESN
KHVDGSGSGSGSGSENLYFQGHHHHHH*

Protein of the Light Chain Fab :
MKYLLPTAAAGLLLLAAQPAMAYELIQPSSASVTVGETVKI
TCSGDQLPKNFAYWFQQKSDKNILLLIYMDNKRPSGIPER
FSGSTSGTTATLTISGAQPEDEAAYYCLSSYGDNNDLVFG
SGTQLTVLRGRTVAAPSVFIFPPSDEQLKSGTASVVCLLN
NFYPREAKVQWKVDNALQSGNSQESVTEQDSKDSTYSL
SSTLTLSKADYEKHKVYACEVTHQGLSSPVTKSFNRGEC*

**Supplementary Figure 1 | Construction of the recombinant LiTe protein.** The LiTe protein is produced in transfected eukaryotic cells following transfection of a plasmid encoding the light chain of the H57-597 Fab and the phytochrome interacting factor 6 (PIF6) domain attached by a flexible linker to the C-terminal amino acid sequence of the heavy chain of the H57-597 Fab. A second linker at the C-terminal part of PIF6 comprised the 6xHis tag for the purification and detection of the recombinant LiTe protein.

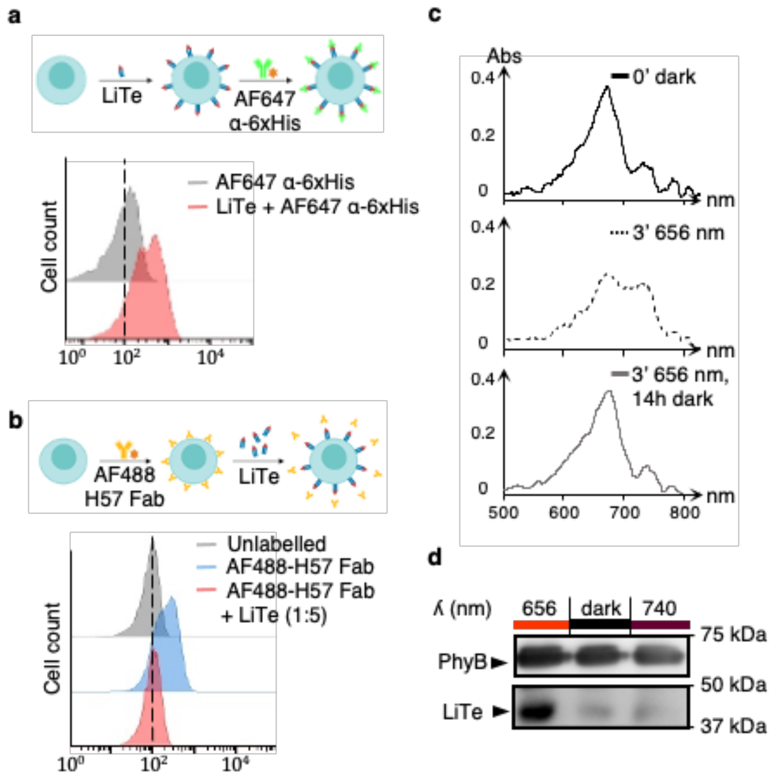


**Supplementary Figure 2 | Evaluation of the functionality of the LiTe system components. a**, Flow cytometry analysis of the binding of LiTe protein to mCD4 T cell line detected by AF647-anti 6xHis mAb. **b**, Displacement of AF488-H57 Fab bound to TCR by the LiTe protein. **c**, Spectral analysis of the PhyB produced in *E. coli*. Top, absorbance of PhyB in a closed conformation, i.e., before 656 nm light illumination. Middle, absorbance spectra in its open conformation induced by a 3 min light exposure at 656 nm. Bottom, same as the middle condition but followed by 14 h incubation in the dark to recover the close conformation of PhyB. **d**, Pull-Down assay of the LiTe protein by PhyB-coated beads co-incubated and exposed to different conditions of illumination (n>3).

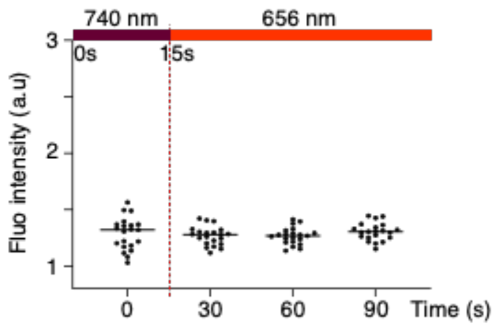


**Supplementary Figure 3 | Light exposure induced no calcium influx in the absence of PhyB.** Primary T lymphocytes were loaded with the PBX calcium sensitive dye and the LiTe protein, and were then maintained in the absence of PhyB-coated beads. Under these conditions, no calcium influxes were observed under 656 nm light exposure (n=20 cells).

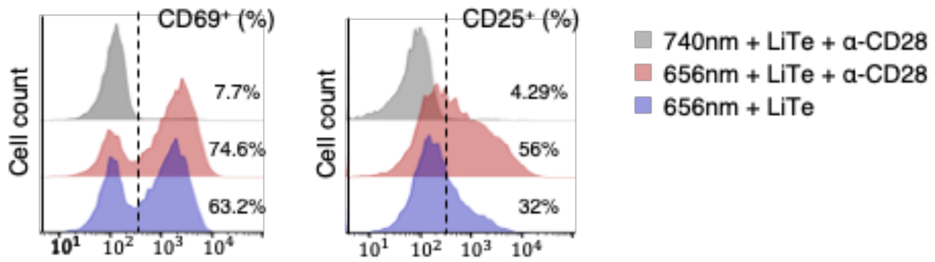


**Supplementary Figure 4 | CD28 costimulation increased the fraction of T cells responding to the LiTe system.** Primary CD8 T cells were incubated with the LiTe system in the presence or not of anti-CD28 antibody, then illuminated 12 h in optoPlate at the specified wavelength. Flow cytometry charts showing the T cell surface expression of CD69 (left) and CD25 (right).

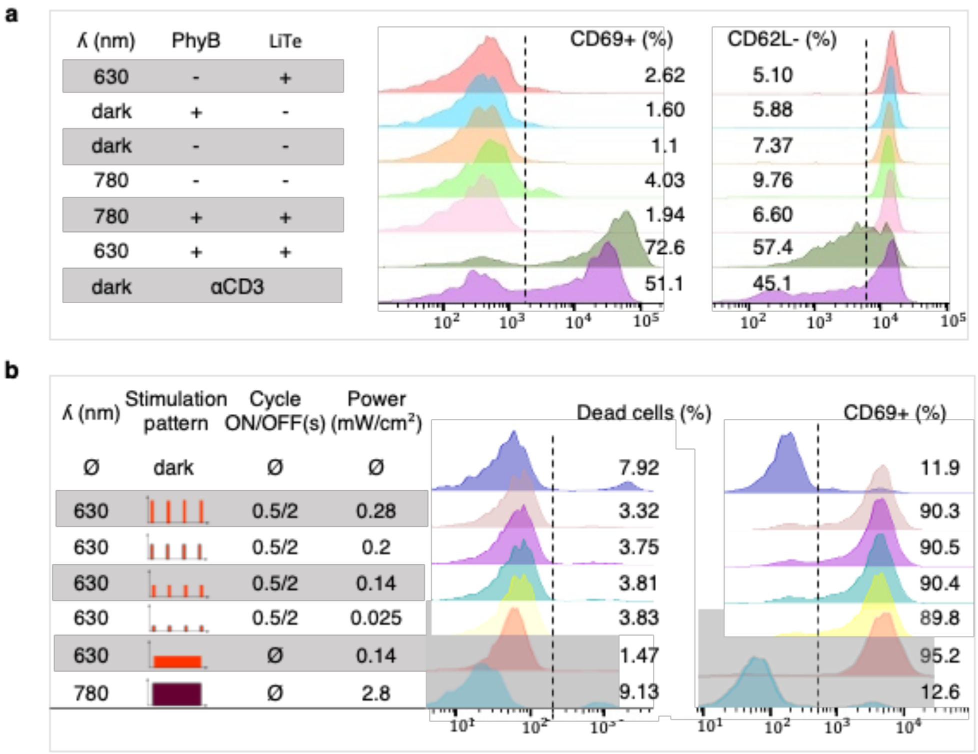


**Supplementary Figure 5 | Control for the LiTe system specificity and for light-induced phototoxicity. a**, Primary T cells were incubated with the LiTe system and anti-CD28 antibody, then illuminated or not for 18 h in the optoPlate at the specified wavelength. Flow cytometry analysis of CD69 and CD62L cell surface expressions in response to 630 or 780 nm light. The activation of T cells required both the LiTe system and red-light exposure. **b**, Same as in **a** but under different illumination conditions. Cell death and T cell activation have been evaluated by flow cytometry using Viobility™ fixable dye and CD69 cell surface expression, respectively. T cell exposure to red or far-red light was not phototoxic.

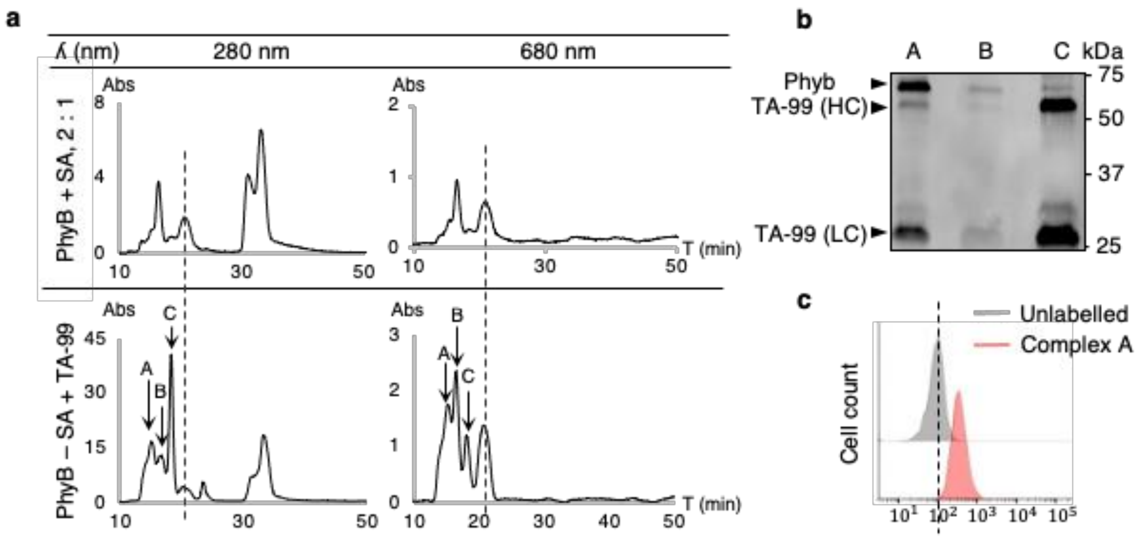


**Supplementary Figure 6 | Production of the Melanoma targeted PhyB.** **a**, Upper panel, complexes of biotinylated PhyB-streptavidin (SA) in 2:1 ratio were analyzed by HPLC followed by absorbance detection at 280 (upper left) and 680 nm (upper right). The dashed line indicates the monomeric form of PhyB. Lower panel, PhyB-SA complexes were incubated with an excess of biotinylated TA-99 mAb and analyzed by HPLC followed by absorbance detection at 280 (lower left) and 680 nm (lower right). **b**, Western blot analysis of the A-C peaks detected by HPLC. A peak corresponds to high molecular weight complexes containing PhyB and TA-99. **c**, Flow cytometry analysis of the A peak complex binding to TRP-1 expressed on B16F10 cells. This A peak complex was used in the LiTe-Me experiments (see Fig. 2 d,e).

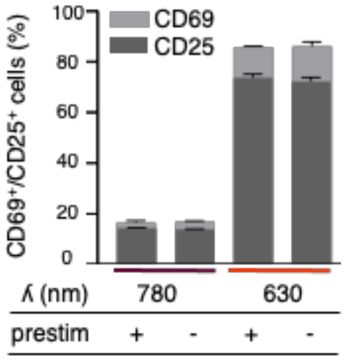


**Supplementary Figure 7 | Conservation of PhyB functionality after iterative 630 nm/780 nm illumination cycles.** PhyB-coated beads were exposed for 4 h to 15 min ON/15 min OFF light cycles (prestim) or not. Then, $CD8^+$ T cells and LiTe were added and the wells exposed to the indicated illumination for 18 h to evaluate by flow cytometry the percentage of CD69 and CD25 positive cells. (n=2; mean +/- SD are shown).
